# Critical assessment of intratumor and low-biomass microbiome using long-read sequencing

**DOI:** 10.64898/2026.02.02.703393

**Authors:** Yanchun Zhang, Edward A. Mead, Mi Ni, Magdalena Ksiezarek, Yujie Liu, Lei Cao, Hao Chen, Yu Fan, Wanjin Qiao, Yangmei Li, Laura Zuluaga, Gintaras Deikus, Robert Sebra, Rachel Brody, Raymund L. Yong, Ketan K. Badani, Xue-Song Zhang, Gang Fang

## Abstract

The detection of low-biomass microbial DNA in human tissues is often confounded by contamination, as demonstrated in the debates over the existence of microbiomes in the placenta, brain, blood, and tumors. Here we show that genomic DNA fragment length serves as a discriminator: while genuine microbiome genomes have long genomic DNA fragments, contaminant DNA is typically short. Using germ-free mouse tissues with bacterial spike-ins and human cell lines, we developed a metric that normalizes microbial read length to host read length. Across multiple human tumor and normal tissues, we found genuine microbiome signals are largely limited to tissues with natural microbial exposure (e.g., gastrointestinal tract, cervix, vagina, skin), while other tissues (e.g. kidney, brain, blood, and placenta) showed no evidence of resident microbiome. These findings support DNA fragment length as a metric for quality controlling low-biomass microbiome profiling, clarifying the debates and strengthen future studies of resident microbiome in tissues.

## Introduction

Low-biomass microbiome samples, such as human tissues, present persistent challenges for metagenomic profiling because of the trace amounts of microbial material they contain(*1*–*4*). Early reports of placental and blood microbiome generated broad interest(*5*–*12*), but subsequent studies revealed that most signals were attributable to contamination from environmental sources or laboratory reagents(*3*, *13*–*18*).

Despite the controversy, the concept of an intratumor microbiome initially appeared highly promising and steadily gained traction, driven in large part by analyses of The Cancer Genome Atlas (TCGA) datasets, which reported cancer-type–specific microbial signatures detectable in tumor tissue and, in some reports, suggesting that microbial patterns could distinguish among many major cancers and complement tumor genomics(*19*–*21*).

However, later re-analyses identified critical oversights, including widespread misclassification of human reads as microbial due to incompleteness or errors in reference genomes and associated databases(*22*–*27*). In one prominent example, millions of apparent bacterial reads detected across TCGA samples were shown to be predominantly human in origin(*23*). This finding challenged both the claims of pervasive tumor–type–specific microbial signals and the nearly-perfect machine-learning cancer classifiers built upon them(*23*). After rigorously eliminating sequences likely attributable to technical artifacts, the residual evidence for tumor-associated microbes appeared far more limited, with the strongest support largely confined to cervical and gastrointestinal cancers(*25*, *27*).

However, some recent studies nonetheless continued to report compelling evidence for intratumor microbial signals even after stringent filtering and contamination-aware workflows(*28*–*31*), highlighting the findings as a solid foundation for a bona fide tumor-associated microbiome. For example, recent work in brain tumors applied multi-modal strategies, including in situ detection of bacterial components, high-resolution imaging, and custom sequencing pipelines, to argue that microbial elements can be detected within the brain tumor microenvironment and that these signals vary by tumor type and anatomical context(*29*, *30*). These studies further proposed that intratumor microbial signatures may correlate with regional immune and immunometabolic programs, raising the possibility that microbial features could help explain heterogeneity in tumor behavior and therapeutic response(*29*, *30*).

Yet the debate remains unsettled, and the mixed literature has created confusion for both microbiome and cancer researchers: it remains unclear whether residual microbial reads reflect genuine endogenous biology, which analytical methods are reliable in low-biomass contexts, and how to reconcile findings across different studies(*1*–*4*). Distinguishing intact microbial cells from fragmented contaminant DNA is essential for establishing reliable and reproducible approaches to studying host-associated microbiota in low-biomass environments.

Here, we used long-read sequencing and built on its ability to preserve genomic DNA (gDNA) fragment length distribution to address this challenge. Using positive and negative controls, we observed that intact microbial cells, reflecting live resident microbiomes, have long gDNA fragments, whereas contaminant DNA is typically short and degraded. Leveraging long-read sequencing, we developed Median Length-Adjusted (*Median(L)adj*), a metric that normalizes microbial read length to host read length, thereby capturing microbial DNA lengths while controlling technical variations. Applying this metric to diverse tumor and normal tissues, we found credible microbial signals enriched in biopsy sites with natural microbial exposure, such as the gastrointestinal (GI) tract, cervix, vagina, and skin. In contrast, tissues traditionally regarded as sterile, including the placenta, brain, blood, and most tumors, showed no evidence of resident microbiota. These findings establish DNA fragment length as a useful quality-control feature for microbiome profiling in low-biomass settings and improve the reliability of future tumor microbiome research.

## Results

### Bacterial gDNA in negative-control tissues and cell lines is short and fragmented

To characterize the nature of bacterial reads persisting after stringent host filtering, we first analyzed germ-free mouse (GFM) tissues, where no live resident gut microbiota are expected. Oxford Nanopore Technology (ONT) long-read sequencing was performed on seven GFM tissue samples from multiple organs (Table S1). After removing host-mapped reads using a rigorously evaluated multi-step pipeline(*23*, *25*)(Fig. 1A; Methods), we observed that bacterial reads remaining in these samples were markedly shorter than mouse gDNA (Fig. 1B). Since host and bacterial DNA were extracted and processed together, the consistent length disparity therefore does not reflect technical degradation. These findings suggest that bacterial sequences in GFM tissues likely stem from degraded DNA contaminants instead of intact genomes from resident cells.

**Figure 1.**
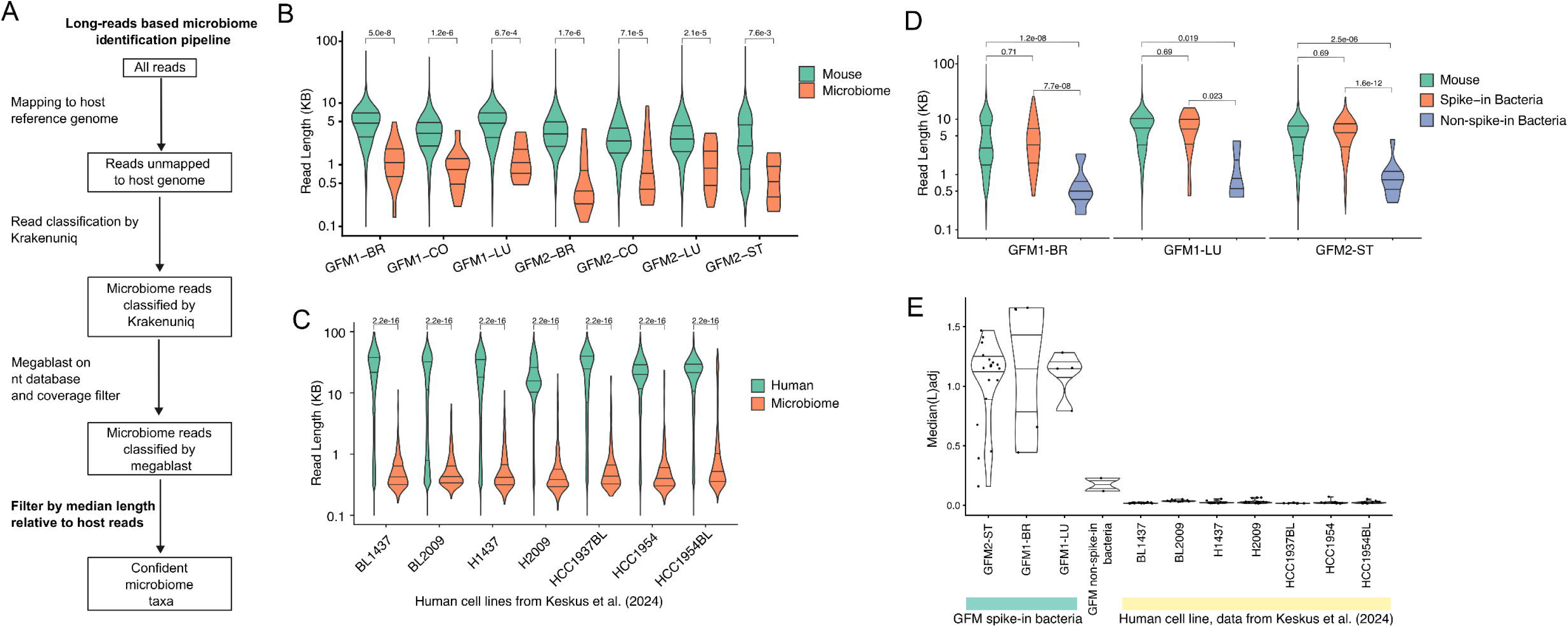
Read-length-based filtering enables reliable microbiome detection in long-read sequencing data. (A) Overview of the computational and analytical workflow used to identify high-confidence microbial taxa from long-read sequencing data. Steps prior to BLAST alignment were adapted from a previously described workflow (*Gihawi et al*., 2023). (B) Read-length distributions of mouse and microbial reads from germ-free mouse (GFM) tissues (n=7). Microbial reads were identified by the workflow from (A). *P* values were calculated by Wilcoxon rank-sum test. (C) Read-length distributions of human and microbial reads in public ONT human cell line datasets (*Keskus et al., 2025*; n=7), analyzed using the same filtering pipeline. *P* values were calculated by Wilcoxon rank-sum test. (D) Read-length distributions of mouse, spike-in bacterial, and non-spike-in bacterial reads from GFM tissues with defined bacteria cell spike-ins (n=3). Spike-in bacteria reads were identified by mapping to the exact reference genome of the spike-in species, and non-spike-in bacterial reads were identified from the remaining unmapped reads using the workflow in (A). *P* values were calculated by Wilcoxon rank-sum test. (E) Median Length-Adjusted (*Median(L)adj*) values of detected spike-in species and non-spike-in bacteria genera in GFM samples and genera from the ONT data of seven public human cell line samples (*Keskus et al., 2025*). Abbreviations: GFM, germ-free mice; BR, brain; LU, lung; ST, stomach.

We next asked whether similar patterns appeared in another negative-control context: cultured human cell lines, which are not expected to harbor live resident microbiota. We re-analyzed matched datasets of seven cell lines sequenced on ONT long-read and matched Illumina short-read platforms(*32*) (Table S2). After applying the same rigorous host–bacteria filtering pipeline (Methods), ONT data again showed that bacterial reads were significantly shorter than human reads across all seven cell lines (Fig. 1C), consistent with the pattern of GFM samples that most microbial reads are fragmented.

Next, to directly test whether intact microbes in host tissue samples yield long gDNA fragments, we introduced bacterial cell spike-ins from 20 species with defined genomes into three GFM tissue and then performed gDNA extraction (Methods). Indeed, spike-in taxa produced long bacterial reads comparable in length to mouse gDNA (median ∼5 kb; Fig. 1D), in sharp contrast to the markedly shorter non–spike-in bacterial reads detected in the same samples. These results confirm that gDNA in intact microbial cells are different from the trace amount of microbial reads in negative controls based on gDNA length captured by long read sequencing.

### A read-length-based metric separates genuine microbiota from contaminants

We hypothesized that the relative lengths of bacterial versus host reads can provide an informative discriminator between intact bacteria and fragmented contaminants. Because read-length distributions vary due to sample storage, DNA extraction protocols and batches (Fig. S1A), we took a normalized approach and developed Median Length-Adjusted (*Median(L)adj*), defined as the ratio of median bacterial read length to median host read length within the same sample (Methods). This metric accounts for variability in sample collection, storage, DNA extraction, and library preparation while retaining the biologically informative signal of bacterial gDNA fragment size relative to host gDNA.

A key premise underlying our framework is that intact microbial cells present at meaningful levels should yield gDNA fragments comparable in length to host DNA when processed under identical extraction and library conditions. While host and microbial DNA may differ in lysis susceptibility, genuine microbial genomes from intact cells should produce a substantial fraction of long fragments overlapping the length scale of host-derived DNA. In contrast, microbial reads arising from degraded environmental DNA, reagent contamination, or other low-integrity sources are expected to be predominantly short.

Applying *Median(L)adj* to spike-in bacteria in GFM tissues and to bacterial genera detected in human cell lines revealed a clear difference: spike-in bacterial reads consistently showed high values, whereas non–spike-in taxa in GFM tissues and cell line bacteria had very low values (Fig. 1E, Table S3). These results demonstrate that *Median(L)adj* provides an effective metric to differentiate genuine resident microbiome taxa from fragmented contaminants in long-read sequencing data.

### ***Median(L)adj*** shows that genuine microbial signals in tumor and normal tissues are confined to GI tissues and other exposure-associated biopsy sites

We next applied *Median(L)adj* to a collection of public long-read sequencing data of multiple types of human primary tumors and normal tissues (249 samples). In addition, we generated ONT sequencing data from six colorectal cancer (CRC) tumors and eight matched normal tissues, five glioma tumors, eighteen kidney cancer tumors and four gastric biopsy samples (Table S4).

We first analyzed the new data we generated along with two public datasets, one from *Xu et al., 2023*(*33*), comprising 21 matched CRC tumor–normal pairs sequenced using ONT (Table S5), and another one from a lung cancer dataset from *Sakamoto et al., 2022*(*34*), which sequenced 23 lung cancer-normal pairs using both ONT and Illumina platforms (Table S6). Across these cancer and normal tissue samples we examined, only CRC and gastric biopsy samples contained bacterial genera with relatively high *Median(L)adj* values (Fig. 2A, S2A, Methods). Specifically, the tumor-associated bacteria genus *Fusobacterium* in colorectal cancer(*35*, *36*) and *Helicobacter pylori* in gastric biopsy samples were detected with relatively high *Median(L)adj* values (Fig. S2B), which are consistent with their well-established genuine association with CRC and gastric tissues, respectively. By contrast, glioma, kidney cancer and lung cancer samples lacked high *Median(L)adj* bacterial genera: because many of them have bacterial read counts too low (less than five reads) to reliably estimate median, we performed an aggregate analyses in which the 18 kidney cancer samples and 46 lung cancer samples were pooled into two pseudo-samples with substantially increased effective depth. Even after pooling, most detected genera continued to exhibit very low *Median(L)adj* values, suggesting that insufficient depth is unlikely to account for the absence of confident microbial signals and that no strong evidence for viable tissue-resident microbes in kidney and lung tissues (Fig. 2A, inset). The microbial signal we observed in lung tissues was characterized by very low biomass and predominantly short, fragmented reads, suggesting that most detected bacterial content is degraded gDNA from transient exposure from breathe or technical carryover, as opposed to resident microbial genomes (Fig. 2B).

**Figure 2.**
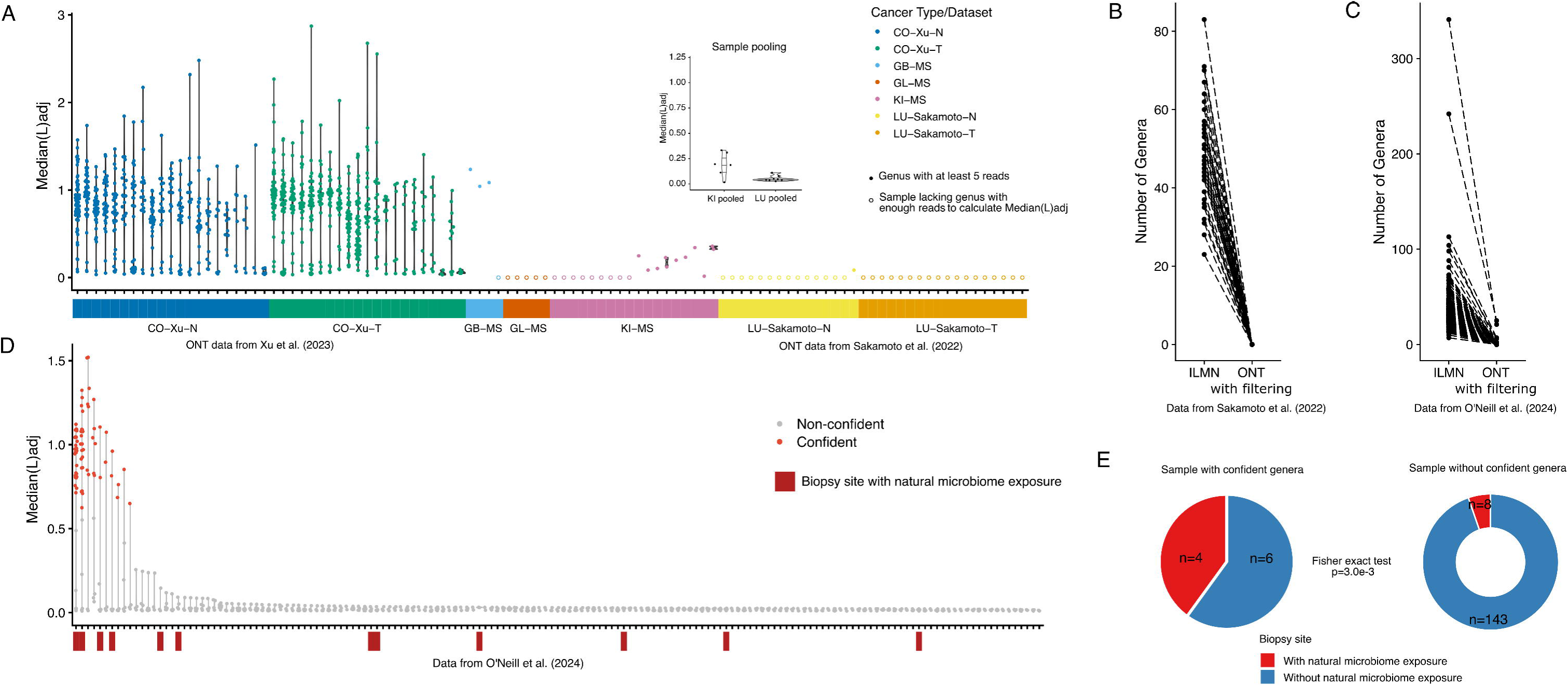
Long-read-based microbiome identification across human cancer and normal tissues. (A) *Median(L)adj* values of detected genera in ONT datasets across multiple human tissue and tumor types. Data includes public CRC tumor (CO-Xu-T; n=21) and matched normal (CO-Xu-N; n=21) samples (*Xu et al., 2023*), public lung cancer tumor (LU-Sakamoto-T; n=23) and matched normal (LU-Sakamoto-N; n=23) samples (*Sakamoto et al., 2022)*, and in-house sequenced gastric biopsy (GB-MS; n=4), kidney cancer (KI-MS; n=18) and glioma (GL-MS; n=5) samples. The 18 kidney samples and 46 lung cancer samples were also pooled into pseudo-samples to assess sequencing-depth effects, with pooled results shown in the upper right inset. Previously reported common reagent-associated contaminant genera (*Salter et al., 2014*) were excluded. Samples lacking sufficient reads for *Median(L)adj* calculation are shown as open circles. (B) Number of genera detected in matched Illumina and ONT datasets from the public lung cancer cohort (*Sakamoto et al., 2022*; n=23 pairs), after applying a *Median(L)adj* cutoff of 0.6 on the ONT data. (C) Number of genera detected in matched Illumina and ONT datasets from a multi-cancer cohort (*O’Neill et al., 2024*; n=161 samples), after applying a *Median(L)adj* cutoff of 0.6 on the ONT data. (D) *Median(L)adj* values of detected genera in ONT data from the multi-cancer cohort (*O’Neill et al., 2024*). Genera with *Median(L)adj* > 0.6 were defined as confident (red) and non-confident genera are shown in gray. Samples from biopsy sites with natural microbiome exposure (e.g., skin, tongue, vagina and cervix) were highlighted in dark red. (E) Proportion of cancer samples originating from biopsy sites with and without natural microbiome exposure in the ONT data from the pan-cancer dataset (*O’Neill et al., 2024*). *P value* of the enrichment of samples with confident genera and biopsy sites with natural microbiome exposure was assessed by Fisher’s exact test. Abbreviations: CO, colon cancer; GB, gastric biopsy; GL, glioma; KI, kidney cancer; LU, lung cancer; N, normal; T, tumor.

To further assess the prevalence of confident microbial signals across cancer types, we reanalyzed a public pan-cancer ONT dataset comprising 161 tumor samples (123 metastatic tumors and 38 local tumors) from 25 cancer types and 36 different biopsy sites (*O’Neill et al., 2024*(*37*)) (Table S7). Again, the ONT data with *Median(L)adj* filtering identified substantially fewer confident bacterial genera than Illumina data (Fig. 2C). Only ten tumor samples (five local tumor and five metastatic) harbored confident microbial signals, and just four contained more than three high *Median(L)adj* genera (Fig. 2D), indicating no widespread presence of intra-tumor microbiome in these samples (Table S8). Four of these samples originated from biopsy sites with natural microbial exposure (e.g., cervix, skin, vagina and tongue), demonstrating strong enrichment for such sites (Fig. 2E). The other six samples, although not categorized into sites with natural microbial exposure, were also from sites that possibly had contact with microbiomes, such as lymph node and chest wall (Table S7, S8). Looking into all the 123 metastatic tumor samples, we observed no association between confident microbial signals and the original tumor type, for example, neither CRC nor melanoma-derived metastases harbored confident microbial taxa (Fig. S3).

### Residual long microbial reads are dominated by well-known contaminants

While *Median(L)adj* provides a genus-level, host-normalized summary of microbial DNA integrity and indicates that tissues without natural microbial exposure lack evidence for prevalent tissue-resident microbiomes, an important follow-up question is: what explains the small number of residual microbial reads that are themselves long? To address this, we performed a targeted taxonomic analysis focusing on microbial reads longer than 5 kb in the human samples analyzed above. In contrast to *Median(L)adj*, which is explicitly designed to normalize microbial fragment lengths to host DNA within each sample, the purpose of this hard-length cutoff is not to make a binary call about “presence/absence” of a microbiome, but to enrich for high-integrity microbial fragments that are least consistent with degraded environmental/reagent DNA and therefore most informative when searching for any credible evidence of resident microbes in otherwise low-biomass contexts.

This long-fragment–focused analysis yielded several observations. First, gastrointestinal (GI) tissues showed abundant microbial reads >5 kb spanning many genera, broadly reflecting the expected diversity of the GI microbiome (Fig. 3A). Second, within GI-associated cancers and other cancers with natural bacteria exposure (e.g., from skin or vagina, et al.), we observed biologically coherent and recurrent signals. For example, *Fusobacterium*, a genus frequently associated with CRC, was widely supported by long reads >5 kb (Fig. 3A, B). In contrast, outside GI and other naturally microbe-exposed contexts, the taxa supported by microbial reads >5 kb were far less abundant and were dominated by organisms commonly reported as laboratory or reagent-associated contaminants (Fig. 3A). For example, in one published cell-line sample we found that essentially all microbial reads >5 kb mapped to *Delftia*, a well-known contaminant in cell culture and laboratory workflows (*38*), consistent with true exogenous introduction rather than endogenous tissue colonization. In the public lung cancer dataset, only one sample contained more than five reads longer than 5 kb, all of which were assigned to *Bradyrhyzobium*, another common reagent-associated contaminant(*38*).

**Figure 3.**
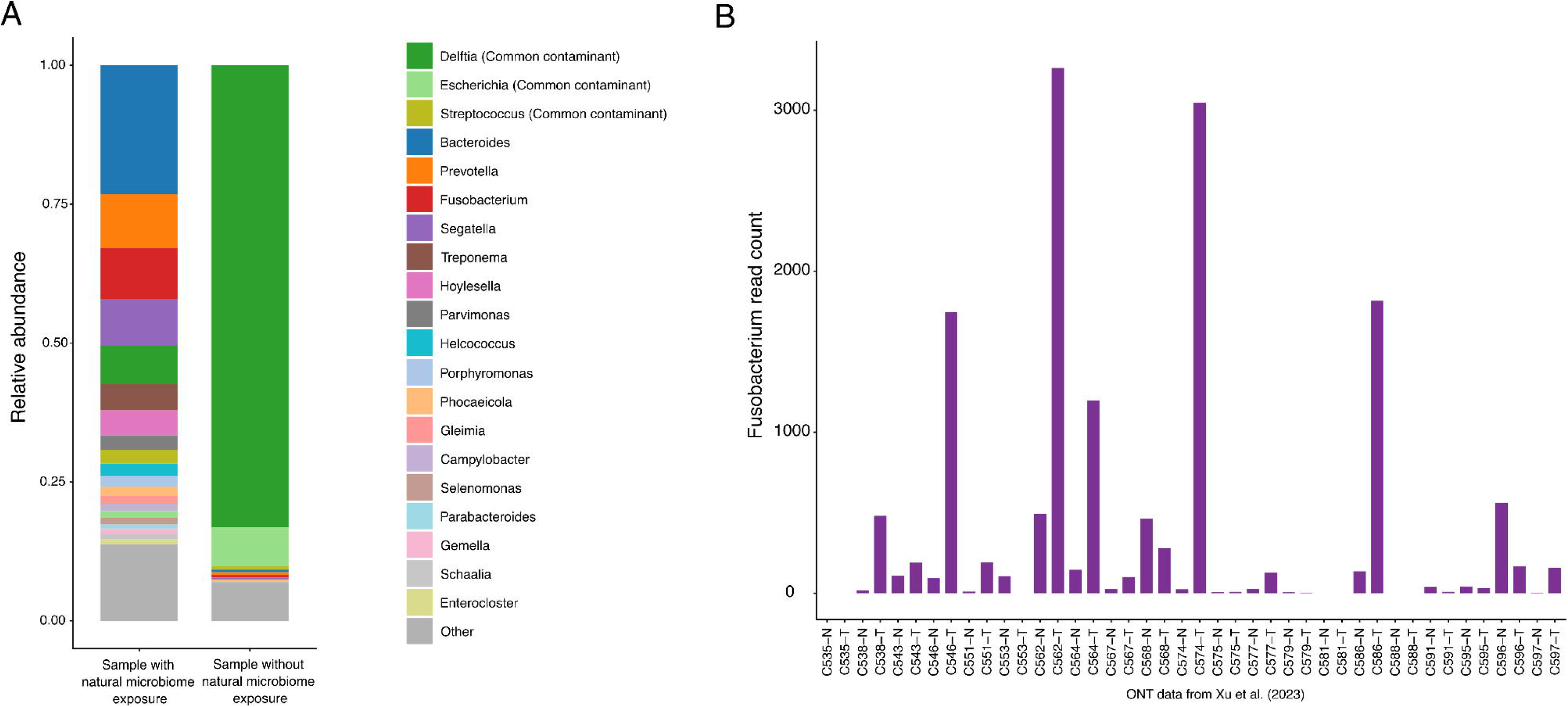
Microbial profiling based on long bacterial reads. (A) Microbial profiles constructed using genus with at least 5 reads longer than 5 kb. Profiles are shown for samples with and without natural microbial exposure. All colorectal cancer samples (*Xu et al., 2023*; n =21 pairs) and samples from vagina, skin, cervix and tongue in the pan-cancer dataset (n=12) (*O’Neill et al., 2024*) were grouped as with natural microbiome exposure, whereas all other samples (n=149) were grouped as without natural microbiome exposure. Genera previously reported as common reagent-associated contaminants (Salter et al., 2014) are labeled as *Common contaminant*. (B) Number of *Fusobacterium* reads longer than 5 kb detected in each sample of the public colorectal cancer dataset (*Xu et al., 2023*; n=42), following the same filtering and length-based criteria.

This targeted residual analysis complements the *Median(L)adj* results by showing that when long microbial fragments are present, they either (i) recapitulate well-established biology in microbe-exposed tissues (e.g. *Fusobacterium* in the gut and CRC) or (ii) are disproportionately explained by well-known laboratory/environmental contaminants in tissues where stable resident microbiomes are not expected. Thus, across the diverse non-microbe-exposed cancer types and sample contexts examined here, we interpret both short and long microbial reads in these tissues as background signal or transient exposure, and find no evidence for a pervasive intratumor microbiome.

### Long-read analysis re-confirms absence of confident microbiome signal in placenta and blood

We then applied our framework to the placenta and blood: two tissues where ‘resident microbiome’ claims have been extensively debated. Current consensus attributes most signals in these tissues to contamination or active infection rather than stable resident microbiota under healthy conditions, aside from bona fide pathogenic infections or severe barrier disruption(*3*, *5*–*7*, *13*–*18*). We therefore examined microbial read-length patterns in placenta and blood as stringent “negative-control-like” tissues to test whether *Median(L)adj* consistently classifies residual microbial reads as fragmented and low-integrity in settings where a resident microbiome is not expected.

We first reanalyzed a public long-read PacBio placenta dataset (*Yu et al., 2021*(*39*), Table S9), a tissue central to the long-standing controversy regarding the existence of a placenta microbiome. Whereas early studies reported microbial signals in the placenta(*5*–*7*), later work largely attributed these findings to contamination(*3*, *13*–*17*). Consistent with this latter view, our analysis detected only short microbial reads and no reliable long fragments (Fig. 4A), providing no evidence for a bona fide placenta microbiome.

**Figure 4.**
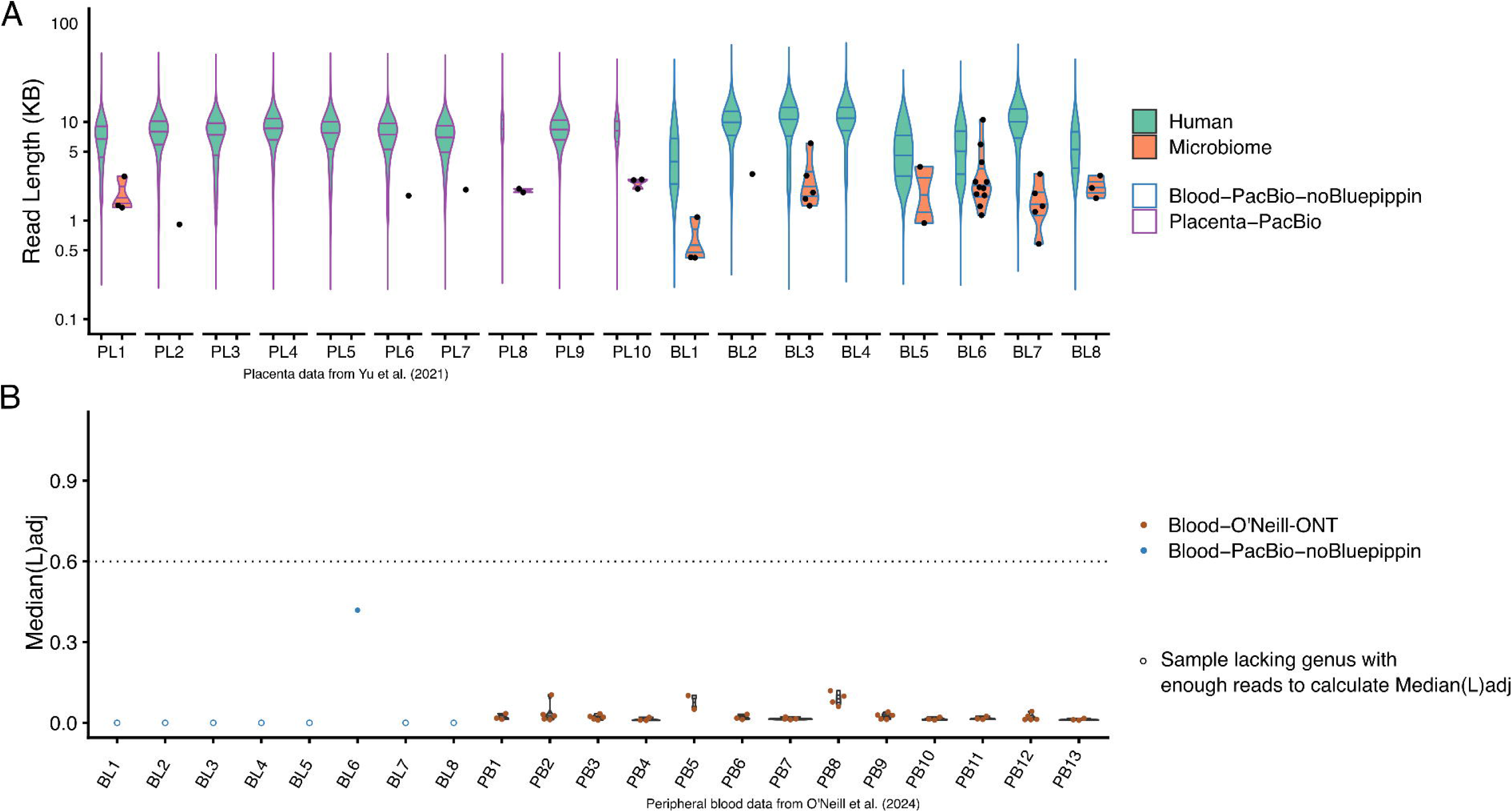
Long-read-based microbial analysis in placenta and blood. (A) Read length distributions of human and microbial reads in public PacBio data of placenta (*Yu et al., 2021*; n=10 samples) and in-house PacBio data of human blood samples (n=8). Blood libraries were prepared without Bluepippin-based size selection. Placenta samples are shown as purple and blood samples in blue. Individual microbial reads are overlaid as dots and each dot represents one read. (B) *Median(L)adj* values of in-house PacBio data of blood sample (n=8) and public peripheral blood ONT data (n=13) (*O’Neill et al., 2024*). In-house samples are shown in blue and public peripheral blood samples in brown. Samples lacking sufficient reads for *Median(L)adj* calculation are shown as open circles. A cutoff of 0.6 was applied. Abbreviations: BL, blood; PL, placenta; PB, peripheral blood from tumor patients (*O’Neill et al., 2024*).

We next generated PacBio long-read data from eight human blood samples without stringent BluePippin-based size selection. Microbial reads were rare and markedly shorter than human reads across all samples (Table S9, Fig. 4A). When examining *Median(L)adj* values in the detected genera in the sequenced blood data and public blood ONT sequencing data from O’Neill’s cancer dataset(*37*), uniformly low values were observed across both datasets (Fig. 4B).

## Discussion

Distinguishing genuine microbiome signals from contamination is a central challenge in low-biomass microbiome research. Using long-reads sequencing, we examined microbial DNA fragment-length distributions as an indicator of genome integrity. Resident microbiota are represented by a majority of intact bacterial cells that yield long DNA fragments, whereas contaminating or degraded material is typically represented by short DNA fragments. We developed the Median Length–Adjusted (*Median(L)adj*) metric, which normalizes bacterial read length to host DNA within the same sample, as a practically useful metric for quality control of genuine microbial taxa across diverse tissues.

Evaluated using rigorously designed negative controls, *Median(L)adj* robustly separated intact bacteria from fragmented contaminants. By contrast, short-read sequencing fragments all DNA molecules into similar lengths, precluding discrimination between intact genomes and degraded material. As a result, reliance on short-read abundance alone can inflate false-positive microbial signals in low-biomass settings, which could be alleviated by our read-length based metric. Consistently, analysis of the “negative-control-like” tissues, including placenta and blood, revealed no evidence of stable resident microbiomes under normal conditions, in agreement with the prevailing view that signals reflect contamination except during active infection(*13*, *18*).

Applying the framework to human tumors and normal tissues, our study showed that in both primary and metastatic cancer, confident microbial signals were largely confined to GI tissues and other sites with natural exposure to microbes, such as skin and vagina. We found no solid evidence for widespread microbiomes across most other tumor types, in contrast to previous short-read–based reports in primary(*19*) and metastatic cancer(*28*). These findings are consistent with the recent The Cancer Genome Atlas (TCGA) reanalysis reporting limited microbiome associations with a small subset of cancers, primarily those from mucosal or other exposure-associated biopsy sites(*25*–*27*). Since these analyses are largely based on short-read sequencing, our long-read-based approach could further reduce false positive microbial signals. Analysis of the metastatic tumor in the public pan-cancer dataset(*37*) suggested that the biopsy site, not the tumor’s tissue of origin, primarily determines whether a confident microbial signal is present. While the earlier short-read–based study reached a similar conclusion(*28*), our findings provide a complementary and clearer evidence.

Reconciling these results with the recent multimodal reports of microbial signals in brain tumors is important(*29*, *30*). A central challenge is that each commonly used modality has distinct limitations in low-biomass contexts, and none alone is sufficient to establish a prevalent, tissue-resident microbiome. For example, 16S rRNA sequencing is highly sensitive but susceptible to reagent and laboratory contamination and provides no information on genome integrity(*40*). Imaging approaches may be confounded by nonspecific staining or residual microbial material introduced during tissue processing that doesn’t reflect living bacteria(*30*). Culture-based assays may preferentially recover exogenous organisms while failing to culture fastidious taxa(*41*). These considerations do not necessarily imply that recent studies are incorrect; rather, they highlight why evidence streams can be individually compelling yet still not conclusive with respect to prevalence, residency, or biological relevance without an integrated framework that explicitly addresses contamination risk and reproducibility. In this context, our long-read-based profiling provides an independent, integrity-based measure for evaluating tissue residency, thereby addressing an ambiguity that abundance- or marker-based approaches cannot resolve. Future studies should therefore evaluate genome-level DNA fragment lengths while integrating multiple technologies under harmonized controls.

The absence of genera with high-integrity bacterial signals in lung tissues was notable. Although the lung is continuously exposed to inhaled environmental microbes and microbial signals have been reported in both lung cancer and adjacent normal tissues(*42*–*47*), the respiratory tract is also equipped with strong barrier and clearance mechanisms, including mucociliary transport, antimicrobial peptides, and vigilant innate immune surveillance(*48*), that tend to prevent stable colonization of living bacteria in the lower airways under healthy conditions(*49*). Deep lung tissue may resemble other low-biomass compartments, such as blood, where bacterial DNA may be intermittently detectable without sustained microbial residency.

Several limitations should be acknowledged. First, the sensitivity of long-read sequencing is constrained by practical sequencing depth, and extremely low-abundance taxa may escape detection. Although our aggregated analyses suggest limited depth does not materially affect our conclusions (Fig. 2A), rare or context-specific colonization events might be detected in future studies with larger cohorts and deeper sequencing. Second, DNA introduced through contamination or transient exposure might occasionally generate long fragments and should not be interpreted, by itself, as definitive evidence of tissue residency.

We would also like to highlight some important considerations to use to apply this read length-based metric in future studies. First, for the clearest distinction between intact and degraded genomes, our long-read analyses in this study were performed on fresh-frozen samples without fixation, which best preserve high-molecular-weight DNA and the native read-length distributions. In contrast, formalin-fixed, paraffin-embedded (FFPE) specimens are generally not recommended for this approach because they typically contain substantial DNA fragmentation, which compresses read-length profiles and makes it difficult to distinguish intact microbial genomes from degraded DNA. Second, library preparation protocols can influence read-length distributions and need to be carefully considered when interpreting long-read data and applying length-based QC. PacBio sequencing commonly incorporates with strict size- selection steps (e.g., BluePippin or equivalent) that substantially deplete short DNA fragments prior to sequencing and consequently reduce sensitivity to fragmented microbial signals(*50*). For ONT long-read sequencing, read length can be protocol-dependent: ligation-based preparations generally preserve high-molecular-weight input DNA and are well suited for integrity comparisons, whereas transposase-based kits can fragment DNA during tagmentation and yield shorter read-length profiles even when sequencing is performed on a long-read platform(*51*). These technical parameters have been accounted for when reanalyzing public datasets or designing new experiments, and key library-prep details (e.g., size selection, ligation vs transposase-based prep, and DNA handling/shearing conditions) should be reported in future work to ensure that read-length–based interpretations remain valid and comparable across studies.

In summary, our study establishes a DNA fragment length–based approach for more rigorous interpretation of microbial signals in low-biomass tissues. Analyses of microbial DNA fragment-length distributions clarifies the longstanding debates surrounding tumor-associated microbiomes and related low-biomass systems. Our results indicate that microbial communities in human tissues are largely confined to exposed anatomical sites and are not a pervasive feature of most primary or metastatic tumors. We encourage future intratumor and other low-biomass microbiome studies to incorporate long-read sequencing and length-aware quality control.

## Methods

### Preparation and DNA extraction of mouse samples

Germ-Free mice (GFM) were bred and maintained at the Rutgers University Gnotobiotic Core facility (Rutgers, The State University of New Jersey). Two germ-free, 8-week-old male C57BL/6J mice were sacrificed, and multiple tissues were collected, including brain, lung, stomach and colon (proximal). All procedures were performed under stringent aseptic conditions to maintain sterility to the greatest extent possible. The gnotobiotic cages (NKP Isotec, Selbitz, Germany) housing each mouse underwent a thorough surface cleaning with detergent solution and then were placed within a secondary sterile hood. The mice were euthanized and dissected using sterilized instruments within a sterile hood. Between each dissection, tools and hood were re-sterilized by a 30-minute UV sterilization cycle. All collected samples were flash frozen in liquid nitrogen and then stored in a -80°C freezer.

Using the dissected mice tissue samples, brain and lung from #1 mice and stomach from the #2 mice were spiked with a mock bacterial mixture consisting of 20 equally abundant strains (MSA-2002; ATCC) onto each tissue sample prior to DNA isolation. For bacterial spike-ins, we used 3.5x10^5^ mock bacteria (“1x concentration”) for the stomach tissue, and 3.5x10^4^ mock bacteria each (“0.1x concentration”) for the lung and brain tissue.

Genomic DNA was isolated by Wizard Genomic DNA Purification Kit (Promega, Madison, WI, USA). Tissues were processed by the animal tissue sub-procedure (3D) of the manufacturer’s protocol. Samples were of varied sizes, but typically 50-250 mg of non-brain samples, and up to 500 mg brain tissue was used. The optional RNase treatment was included. For large samples, scale up of the process was done using several volumes at each step, and typically yielding more DNA at the end. Final elution was carried out in 0.1x IDTE (IDT, Coralville, IA, USA), a 1:9 dilution of the TE stock in Nuclease-Free (NF) Ultrapure water (ThermoFisher Scientific, Waltham, MA, USA).

### Collection and processing of human tissue and blood samples

Colorectal cancer samples were obtained from the Biorepository and Pathology Core at the Icahn School of Medicine at Mount Sinai. Tissue samples were categorized as either tumor core or adjacent normal tissue, with the distance from the tumor (1-9 cm) noted at the time of collection. This spatial gradient was used to model the samples as being gradually more ‘normal’ (non-malignant) as the distance from the tumor increased. Samples had been flash frozen on liquid nitrogen, then stored at -80°C until gDNA isolation by Wizard kit as detailed above.

Glioma and kidney cancer samples of varied sizes were obtained flash frozen from surgeries conducted by the Departments of Neurosurgery and Urology at The Mt. Sinai Hospital (MSH), New York, NY, USA. These had been carefully sectioned by the Biorepository and Pathology CoRE at Icahn School of Medicine at Mount Sinai and provided to each surgeon’s lab, who shared them for DNA isolation. Histopathological assessment confirmed that the glioma samples were all Glioblastoma Multiforme (GBM) samples from various brain regions, while kidney cancers were likewise classified, and most were Renal Cell Carcinomas (RCC; Table S4). The samples were further dissected to achieve distinct pieces from adjacent normal, periphery, and tumor core material, for clinical and research use. Samples had been flash frozen on liquid nitrogen, then stored at -80°C until gDNA isolation by Wizard kit as detailed above for mouse samples.

Gastric biopsy samples were collected from stomach of *H. Pylori* positive patients from Germany(*52*). Sample summary and medical status are listed in the Table S4 of which three were normal and one was gastric adenocarcinoma. The biopsies were cut into tiny pieces with a sterilized blade, then DNA extraction was performed with the DNeasy Blood and Tissue kit (Qiagen #69506) following the tissue DNA extraction protocol.

Whole blood samples of approximately 3-10 mL were taken in purple vacutainers (EDTA) either before brain surgery, or before and immediately after surgery; they were immediately processed. Blood processing prior to gDNA isolation involved spinning down at 2000 xg for 10 minutes at 4°C to separate plasma (plasma-depleted blood was immediately flash frozen in liquid nitrogen and stored at -80°C), followed by spinning the plasma at 12,000 xg for 2 min at 4°C to pellet exosomal material. After this, both the pellet and the plasma supernatant were flash frozen.

Aliquots of plasma-depleted blood, typically 1.5 mL each, and glioma samples, ∼50-250 mg in size, were used for gDNA isolation by Wizard Genomic DNA Purification Kit (Promega, Madison, WI, USA). Blood gDNA isolation was processed based on the whole blood sub-procedure (3A) in the manufacturer’s protocol and the optional RNase treatment was included. For larger samples, scale up of the process was done using several volumes at each step. Final elution was carried out in 0.1x IDTE (IDT, Coralville, IA, USA), a 1:9 dilution of the IDTE stock in Nuclease-Free Ultrapure water (NFW) (ThermoFisher Scientific, Waltham, MA, USA).

DNA quantity for all samples above was assessed by a Qubit 4 fluorometer using the 1xdsDNA High Sensitivity kit (ThermoFisher Scientific).

### ONT library preparation and sequencing

For GFM tissue samples without bacteria spike-in, seven tissue samples with high quality mouse DNA extracted were chosen for ONT sequencing.

For gastric samples, long-read libraries were prepared with the Rapid ONT kit (SQK-RPB004, Oxford Nanopore Technologies) following its protocol. Then the pooled library was loaded onto an R9 MiniON flowcell for sequencing.

For all other samples, IZ1μg DNA was sheared for each sample to 10 kb (6000 RPM) with a g-TUBE (Covaris). After Ampure XP bead cleanup, we used Native Barcoding Kit 24 V14 (SQK-NBD114.24) for preparing R10 libraries and loaded the libraries onto either MinION or PromethION cells. Manufacturer’s instructions were followed except as noted. DNA repair/end prep was increased to 30 minutes at 20°C. No concomitant increase of time at 65°C was needed because 5 minutes is sufficient to denature the enzyme. Barcoding and adapter ligations were done for 1 hour each. We increased elution times on beads to 45’ to improve yields, and used Ampure PB beads at the end at 0.37x, a stricter cutoff to reduce the amount of short reads. MinION and PromethION cells were run for 96 hours, unless cells were cut off early due to run error or pores dying off. Washes and reloads were typically conducted each day of the run to maximize output using the Flow Cell Wash Kit XL (EXP-WSH004-XL; Oxford Nanopore Technologies). When possible, the library was re-used in reloads to minimize the total amount of library that needed to be prepared each time. Basecalling was turned off during runs to minimize the risk of crashing.

### PacBio library preparation

DNA was sheared for each sample to 20 kb (4200 RPM) with a g-TUBE. After Ampure bead cleanup, SMRT (PacBio, Menlo Park, CA, USA) libraries were prepared by SMRTbell prep kit 3.0, and run on a Revio instrument. We carried out end repair for 1hr, ligation from 3 hours to overnight, and used longer elution times on beads to 45 minutes at 37C to improve elution. We use Ampure PB beads at the end (0.5x) instead of the diluting procedure indicated, to help with library retention. BluePippin was not applied to the blood samples. The S1 External marker used. This allowed a very sharp cutoff, excluding reads below 8 kb in size. Eluted samples were combined with a Tween-20 rinse of the well with the solution by the manufacturer, and all were cleaned up with Ampure PB beads. Typically, 2 samples were pooled per library prep.

### Sequencing of PacBio library

PacBio sequencing data collection and processing were carried out at Histogenetics (Ossining, NY). Whole-genome sequencing (WGS) was conducted on the PacBio Revio platform according to the manufacturer’s standard workflow for non-SPRQ runs (SMRT Link v25.1). Annealing of SMRTbell libraries with sequencing primers was performed and then the libraries were subsequently bound to DNA polymerase using the Revio Polymerase Kit (PacBio Cat. No. 102-739-100). Following purification, the polymerase-bound SMRTbell complexes were quantified using a Qubit Fluorometer (ThermoFisher Scientific) and diluted to a final concentration of 300 pM. These libraries were combined with the loading buffer and a sequencing control before being loaded onto SMRT Cells 25M. Data were acquired over 30-hour runs for each SMRT Cell under standard run conditions with adaptive loading enabled.

### ONT raw sequencing data processing

For GFM samples, raw pod5 files were basecalled with the basecaller command in dorado (https://github.com/nanoporetech/dorado) v0.5.3 using 5mCG/5hmCG methylation calling and aligned to a customized reference genome comprising mouse mm10 and the genomes of the 20 species in ATCC MSA-2002. The resulting bam files were demultiplexed with the demux command in dorado v0.5.3. Debarcoded bam files were used for downstream microbial read identification.

For sequenced kidney cancer, glioma and colorectal cancer (CRC), basecalling and demultiplexing were performed as above, except reads were aligned to the human reference genome hg38.

### Public ONT and PacBio data download, processing, and mapping

For ONT data from human cell line cultures, SRA files were downloaded using the prefetch command in sratoolkit v3.1.1 and converted to fastq files with fastq-dump.

For the public CRC ONT dataset(*33*), ONT fastq files were downloaded from https://ngdc.cncb.ac.cn/gsa-human/browse/HRA002638. For the lung cancer dataset(*34*), ONT fastq files were obtained from the NBDC Human Database. All fastq files were aligned to hg38 using minimap2 v2.24(*53*, *54*) to generate bam files. For the O’Neill’s pan-cancer ONT dataset(*37*), pre-mapped bam files were downloaded from EGA using pyega3 and used directly.

For the public placenta PacBio dataset(*39*), raw sequencing subreads bam files were provided by the the Chinese University of Hong Kong (CUHK) Circulating Nucleic Acids Research Group (CNARG). Consensus reads were generated from subread files using the ccs command, then aligned to hg38 using pbmm2 v1.1.0. For sequenced blood PacBio data, HiFi bam files were converted to fastq with samtools(*55*) fastq, then aligned to hg38 using minimap2 v2.24. All bam files generated or obtained in this step were used for downstream long-read microbial read identification.

### Public Illumina data download and mapping

For human cell line datasets(*32*), SRA files were downloaded from NCBI with the prefetch command in sratoolkit v3.1.1, converted to fastq with fastq-dump, and aligned to hg38 using bowtie2 v2.2.3(*56*). For the O’Neill’s cancer dataset, pre-mapped bam files (aligned to hg38) were downloaded from EGA using pyega3. All bam files generated or obtained in this step were used for downstream short-read microbial read identification.

### Identifying confident microbiome reads in ONT and PacBio datasets

For each sample, unmapped reads were extracted from bam files with samtools v1.21 and re-aligned to the T2T reference genome CHM13v2.0(*57*) using minimap2 v2.24. Reads that remained unmapped were classified with krakenuniq v1.0.4(*58*) using the MicrobialDB database (https://benlangmead.github.io/aws-indexes/k2).

Candidate microbial reads identified by krakenuniq were validated with the megablast module of blast+ v2.13 using parameters: -outfmt "6 qseqid sseqid evalue pident length qstart qend sstart send bitscore stitle staxids sscinames" -max_target_seqs 5. The ‘nt’ database (version 2024-08-31) was downloaded from https://ftp.ncbi.nlm.nih.gov/blast/db/.

For each read, the best hit was chosen based on highest sequence identity and coverage using a custom script. Only reads with >50% target coverage and longer than 100 bp were retained as reliable microbial reads.

### Identifying confident microbiome reads in Illumina datasets

For Illumina datasets, paired unmapped reads were extracted and re-aligned to the T2T reference genome using bowtie2 v2.2.3. Remaining unmapped reads were classified with krakenuniq v1.0.4 in paired-end mode using the same database. Paired reads were merged into a single sequence with a 20N linker between mates and used as input for blast megablast v2.13 with the same parameters as above. For each merged read, the best hit was determined based on sequence identity and coverage, and only reads with >50% coverage of the best hit was retained.

### Calculation of Median Length-Adjusted (***Median(L)adJ***) in long-read data

To normalize for variability in read length across samples, we calculated a Median Length-Adjusted (*Median(L)adj*) value, defined as:

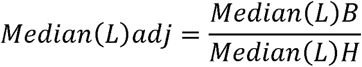

*Median(L)B* represents the median length of bacteria reads. *Median(L)H* represents the median length of host reads. This metric provides a normalized measure of bacterial read length relative to human reads.

For each bacterial genus with ≥5 reads per sample, a *Median(L)adj* value was calculated. Previously reported common reagent contaminant genera (Table S10) were excluded for downstream analysis.

### Categorization of biopsy sites in the O’Neill’s cancer dataset

For samples in the O’Neill’s dataset, biopsy sites containing the keywords “skin”, “tongue”, “vagina” and “cervix” were classified as sites with natural microbiome exposure (hereafter referred to as “exposure_sites”). Samples originating from lymph nodes were excluded from this category.

This classification was used in downstream analyses to assess whether the presence of confident microbial genera (i.e., those with high *Median(L)adj* values) was enriched in exposure-associated biopsy sites compared to non-exposure sites. Fisher’s exact test was used to test the significance of enrichment.

### Long-read profiling of reads longer than 5 kb

Microbial composition profiles were generated from long-read sequencing data by selecting genera supported by at least five reads longer than 5 kb across sequenced kidney cancer and glioma samples, as well as public human cell line, CRC, lung cancer dataset and O’Neill’s pan-cancer datasets. Read lengths were aggregated at the genus level and further grouped by exposure status (with or without natural microbiome exposure). All public CRC samples and samples annotated as exposure-associated sites in the O’Neill dataset were grouped as having natural microbiome exposure. All the remaining samples from the O’Neill’s dataset, the public lung cancer dataset, public human cell lines, and the sequenced glioma and kidney samples were grouped as without natural microbiome exposure. Within each group, genus-level read lengths were summed across samples and converted to relative abundances by normalizing to the total length of long microbial reads in that group.

## Data availability

The sequencing data generated in this study have been deposited in the NCBI Sequence Read Archive under accession number PRJNA1401852. Public human cell lines short-read and long-read sequencing datasets were obtained from NCBI SRA under accession PRJNA1086849. Public colorectal cancer long-read sequencing dataset was obtained from the China National Center for Bioinformation (CNCB), under accession number HRA002638. Controlled-access lung cancer short-read and long-read sequencing datasets were obtained from the Japanese Genotype–Phenotype Archive (JGA) under accessions JGAD000252, JGAD000253, JGAD000463 and JGAD000696 and were accessed with approval from the NBDC Data Access Committee. Controlled-access multiple cancer genomic sequence dataset for long-read and short-read platforms have been obtained at the European Genome-Phenome Archive (EGA; https://ega-archive.org/) as part of study EGA: EGAS00001001159, with approval from the BC Cancer. Controlled-access placenta long-read sequencing dataset was obtained from EGA under accession EGAS00001005515 with approval from the Chinese University of Hong Kong (CUHK) Circulating Nucleic Acids Research Group (CNARG).

## Acknowledgments

This work was supported in part by the staff and resources of the Center for AI-Driven Genomic and Microbiome Medicine and Department of Scientific Computing at the Icahn School of Medicine at Mount Sinai. This work was supported by grant no. R35 GM139655 (G.F.) from the National Institutes of Health. This research was supported by the Biorepository and Pathology Core at the Icahn School of Medicine at Mount Sinai. We thank Dr. Rachel Brody and Pathologist, and the Core laboratory staff and biorepository participants for their invaluable contributions. We are also grateful to the surgeons, Drs. Celia Divino, Spiros Hiotis, and Sergey Khaitov, and pathologists, Drs. Qingqing Liu, Alexandros Polydorides, and Xintong Wang, involved in the care of patients whose samples were included in this study. We would like to acknowledge Kenny Jing Lo, MS, Drs. Indu Saini and Kennedy Okhawere, Jewel Bamby, and Chitra Hindnavis, MA, from Dr. Ketan Badani’s laboratory in Urology at Mount Sinai for their assistance with sample and data collection of kidney samples. We would also like to acknowledge HistoGenetics LLC for their SMRT sequencing. Hwa Ran Kim (Histogenetics) provided additional guidance on optimizing SMRT library preparation. A portion of the data (lung cancer dataset, Sakamoto et al.) was originally obtained by the research group led by Dr. Yutaka Suzuki and Dr. Akayo Suzuki and is available through the NBDC Human Database of the Database Center for Life Science (DBCLS) / the Joint Support-Center for Data Science Research (DS) of the Research Organization of Information and Systems (ROIS) under accession numbers JGAS000349 and JGAS000065. This manuscript was prepared using a limited access dataset obtained from BC CANCER and does not necessarily reflect the opinions or views from BC CANCER. This work would not be possible without the participation of the patients and families, the POG team, the GSC platform, and the generous support of the BC Cancer Foundation, Genome British Columbia (project B20POG, MOH001, MOH002), and the Terry Fox Research Institute Marathon of Hope Cancer Centres Network. This study also utilized data generated by The Chinese University of Hong Kong (CUHK) Circulating Nucleic Acids Research Group, as reported by Chan et al. in *Clinical Chemistry* (doi:10.1373/clinchem.2012.196014).

## Author contributions

G.F. conceived the study and supervised the research. E.A.M., Y.Z., and G.F. designed the experiments. Y.Z. performed all computational analyses. E.A.M. performed most experiments. X.-S.Z. prepared the germ-free mouse samples. Y.Z. and G.F. applied for and curated public sequencing datasets. Y.Liu performed ONT sequencing of gastric biopsy samples. M.N. processed raw ONT data from gastric biopsy samples. L.C. and E.A.M. prepared PacBio libraries from blood samples. G.D. and R.S. advised on PacBio library preparation. R.B. collected colorectal cancer samples. L.Z. and K.K.B. collected kidney cancer samples. R.L.Y. collected glioma samples. Y.Z., E.A.M., and G.F. wrote the manuscript. M.K., M.N., Y.Liu, L.C., H.C., Y.F., W.Q., Y.Li and X.-S.Z. contributed to data interpretation and manuscript editing. All authors reviewed and approved the final manuscript.

## Ethics declarations

All animal procedures were conducted following protocols approved by the Rutgers University Institutional Animal Care and Use Committee (IACUC; protocol numbers 201900013 and 201900032).

Colorectal cancer samples were provided de-identified and as such did not require an IRB.

The kidney cancer samples used were under study IRB #19-00152 that was approved by the Institutional Review Board (IRB) of Mount Sinai. All tumor, blood and urine samples were collected with informed consent from participants in accordance with ethical guidelines. The participants were fully informed about the study’s aims and the voluntary nature of their participation, including the use of their samples for research purposes. Patient confidentiality was maintained throughout the study, and all procedures were conducted in compliance with relevant ethical standards.

The glioblastoma and blood samples were collected under IRB# STUDY-20-01197, approved by the Institutional Review Board (IRB) of Mount Sinai.

The gastric biopsy samples were originally collected in Germany under approval from the local ethics committee, as reported by Venerito in doi:10.1111/j.1365-2036.2006.02856.x.

The authors declare no conflicts of interest related to this study.

