## Supplemental Figure 1-3 for "Critical assessment of intratumor and low-biomass microbiome using long-read sequencing"

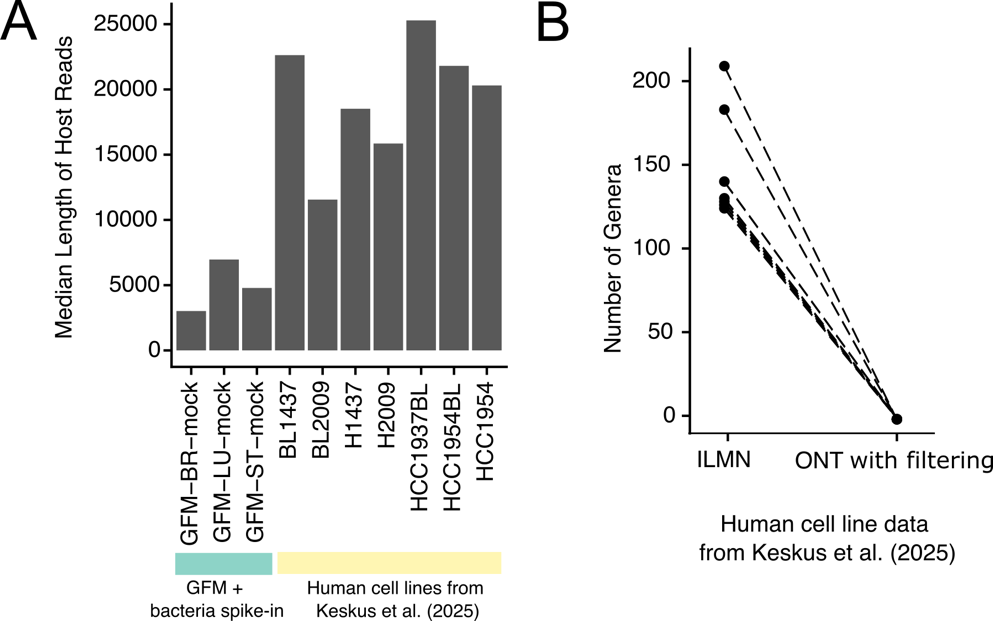

**Figure S1. Read length-based filtering identifies reliable microbial taxa in long-read sequencing data.**

(A) Median lengths of host (mouse/human) reads in germ-free mouse samples and public human cell line samples (*Keskus et al., 2025*), illustrating substantial variation in read-length distributions across samples.

(B) Number of microbial genera detected in ONT data with $Median\left( L \right)adj$-based filtering, compared with matched Illumina datasets from public human cell line samples (*Keskus et al., 2025*). A cut off of 0.6 was applied on the ONT data.

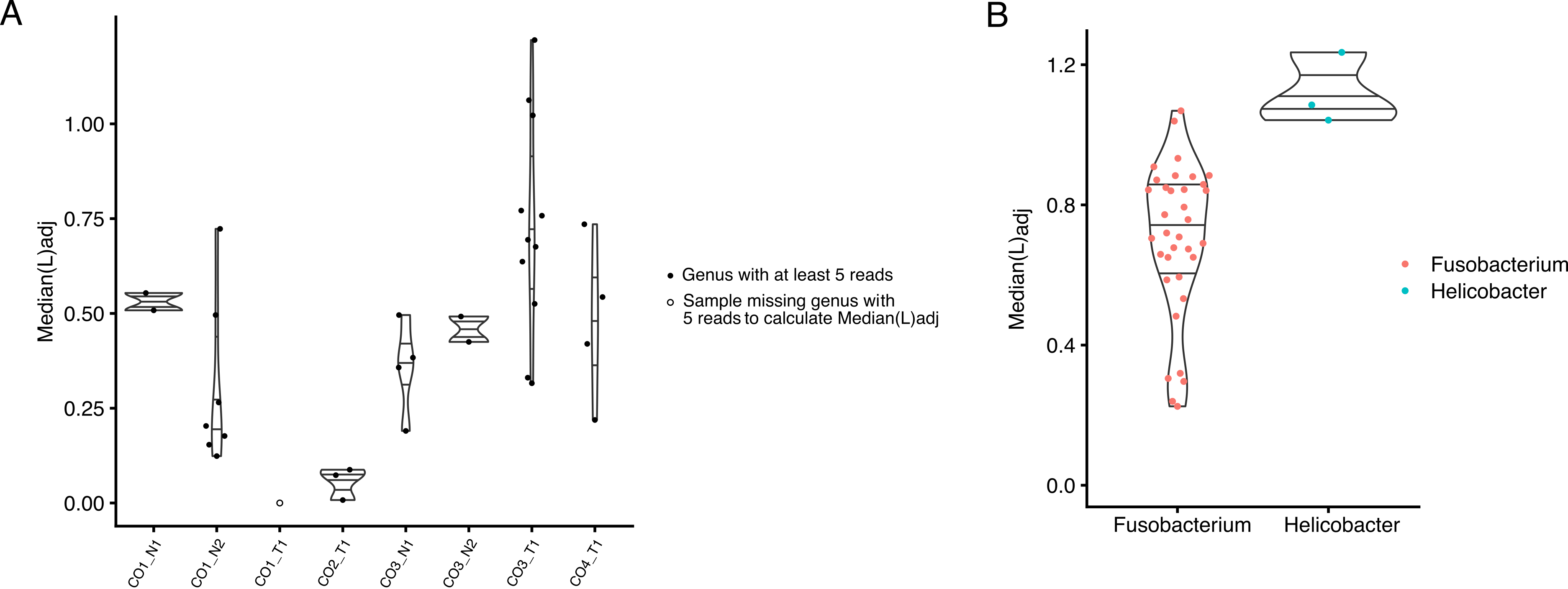

**Figure S2. Long-read-based microbiome identification across tissue and cancer types.**

(A) $Median\left( L \right)adj$ values of detected genera in ONT data from colorectal cancer (CRC) tumors and matched normal tissues sequenced in this study. Samples lacking sufficient reads for $Median\left( L \right)adj$ calculation are shown as open circles.

(B) $Median\left( L \right)adj$ values of *Fusobacterium* from public CRC tumor and matched normal samples (*Xu et al., 2023*) and *Helicobacter* from ONT data of three gastric biopsy samples sequenced in this study.

Abbreviations: CO, colon cancer; N, normal; T, tumor.

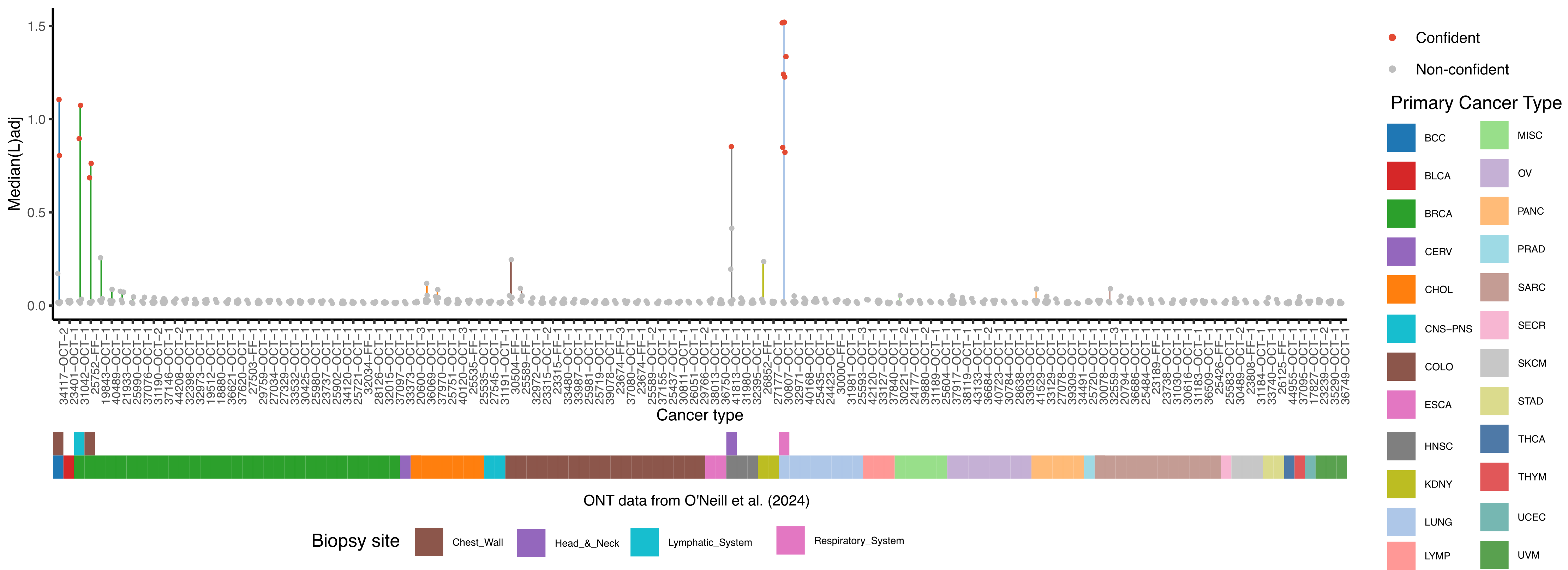

**Figure S3**. $Median\left( L \right)adj$ values of detected genera in the metastatic tumor samples (n=123) from the pan-cancer cohort (*O’Neill et al., 2024*), annotated by primary cancer type and biopsy site. Genera with $Median\left( L \right)adj$ > 0.6 were defined as confident (red) and non-confident genera are shown in gray. Legends for biopsy sites and primary cancer types are shown at the bottom and right, respectively.

Abbreviations: BCC, basal cell carcinoma; BLCA, bladder urothelial carcinoma; BRCA, breast cancer; CERV, cervical cancer; CHOL, cholangiocarcinoma; CNS-PNS, central nervous system and peripheral nervous system cancer; COLO, colorectal cancer; ESCA, esophageal cancer; HNSC, head and neck squamous cell carcinomas; KDNY, kidney cancer; LUNG, lung cancer; LYMP, lymph cancer; MISC, miscellaneous; OV, ovarian cancer; PANC, pancreatic cancer; PRAD, prostate adenocarcinoma; SARC, sarcoma; SECR, adenoid cystic carcinoma of the trachea; SKCM, skin cutaneous melanoma; STAD, stomach adenocarcinoma; THCA, thyroid carcinoma; THYM, thymoma; UCEC, uterine corpus endometrial carcinoma; UVM, uveal melanoma.
